# Internal Grant Review: A Pre-Submission Program for Early-Career Clinical and Translational Researchers

**DOI:** 10.64898/2026.09.21.753152

**Authors:** Amy M. Harrigan, Karen C. Johnston, Jennifer Kirkham, Jason A. Papin

## Abstract

The iTHRIV Scholars Mentored Career Development Program initiated an Internal Grant Review (IGR) program in 2020 for current and recently graduated Scholars seeking funding through K and R awards from the NIH. The IGR program is designed to replicate the NIH review process and provide Scholars the opportunity to receive valuable feedback on their applications prior to NIH submission. A key characteristic of the program is the integration of REDCap, enabling automation and year-round offering while improving tracking and reporting efforts and capabilities. Results of the program have been overall positive, both in proposal development and participant feedback. IGR is a sustainable, valuable resource for early-career faculty competing for limited resources in the pursuit to become independently funded clinical translational scientists.

## Introduction

The odds are against early career researchers pursuing extramural funding through the NIH; the competition for funding grows more rapidly than the number of applicants awarded per year. According to the most recent NIH data, the peak success rate for all competing applications in research grants in the past two decades was at 34% in 1999; in 2025 it was 14%. In this same timeframe, the number of competing applications has more than doubled.^1^ Interestingly, resubmissions and revisions of an application submitted during the same fiscal year as the original application are excluded in these numbers,^2^ meaning it is safe to assume that the current success rate is even less than reported.

To address this and other challenges early career investigators often encounter, the integrated Translational Health Research Institute of Virginia (iTHRIV)^3^ established the iTHRIV Scholars Mentored Career Development Program (iTSP) in 2017. This competitive two-year program is designed to train appointed early career clinical and translational researchers located at iTHRIV partner institutions, the University of Virginia, Virginia Tech, and Carilion Clinic. It is partially funded by the National Center for Advancing Translational Science through the National Institutes of Health Award UL1TR003015/KL2TR003016.

A major goal for all iTHRIV Scholars is to become independent translational investigators, and many, if not all, are pursuing NIH funding to achieve this goal. Recognizing the need for additional support in this area, iTSP initiated the Internal Grant Review (IGR) program in December 2020, an exclusive resource for iTHRIV Scholars and recent iTSP graduates. The original purpose of the IGR program was to provide Scholars with NIH proposals reviewer scores and feedback prior to NIH submission, with two main objectives: 1) to improve the proposal for final submission to the NIH, and 2) to provide Scholars with valuable experience in receiving scores and feedback, as most lack it. These objectives are similar to mock review programs established at other institutions,^4-6^ although there are differences in our approach, such as providing optional discussion of written reviews, assigning every application to a biostatistician reviewer, and allowing Scholar participants flexibility in selecting their deadlines. After offering this resource for several years, we have created infrastructure that drastically improves efficiency and has allowed us to make this a year-round offering. The IGR program fulfills its original purpose and may also foster important lessons in resilience, time management, and the value of seeking diverse perspectives.

### Structure of the Internal Grant Review Program

The IGR program is designed to serve as a pre-NIH review for iTHRIV Scholars (current and recent graduates) applying for NIH K and R awards. Eligible participants in IGR are required to plan ahead during their application preparation to account for the time it takes for reviewing and incorporating revisions prior to NIH submission. Scholars who participate in the IGR program are highly encouraged to submit their full application in near final form to maximize the benefit of a pre-NIH review. If a Scholar chooses to submit a partial application for IGR, there are specific required components they must include for the review, such as the Specific Aims, Research Strategy, and Career Development Plan (for K awards).

Scholars interested in participating in IGR must complete an “Intent to Participate” form in REDCap^7^ which initiates the administrative process of determining deadlines and securing reviewers. Scholars are asked to select a date for their reviews to be returned to them during registration. It is highly encouraged that the Scholars give themselves a month or more to incorporate the reviewer comments into their application revisions, but Scholars can ultimately choose to get reviews back as late as two weeks before NIH submission. Information collected during registration provides the iTSP Leadership with the necessary details to identify and secure appropriate reviewers from our iTHRIV institutions. Reviewers are identified based on their expertise relevant to the application and/or success achieving funding from the NIH award type for which the Scholar is applying. Three to five unpaid reviewers, including a biostatistician, are selected for each application.

Reviewers are given two weeks to review the applications and complete an award-specific (K or R) review form modeled after ones used in NIH study sections. Instructions to reviewers are provided, along with encouragement to reach out to the iTSP Manager for any questions or assistance. Upon submitting their reviews, each reviewer is asked to indicate their willingness to discuss their critiques with the Scholar, if the Scholar wishes. Any reviewer not willing to discuss their review will remain anonymous to the Scholar. All reviews receive an administrative check before being distributed to the Scholar by their chosen due date. More details for the IGR process are detailed in a flowchart in Figure 1.

**Figure 1.**
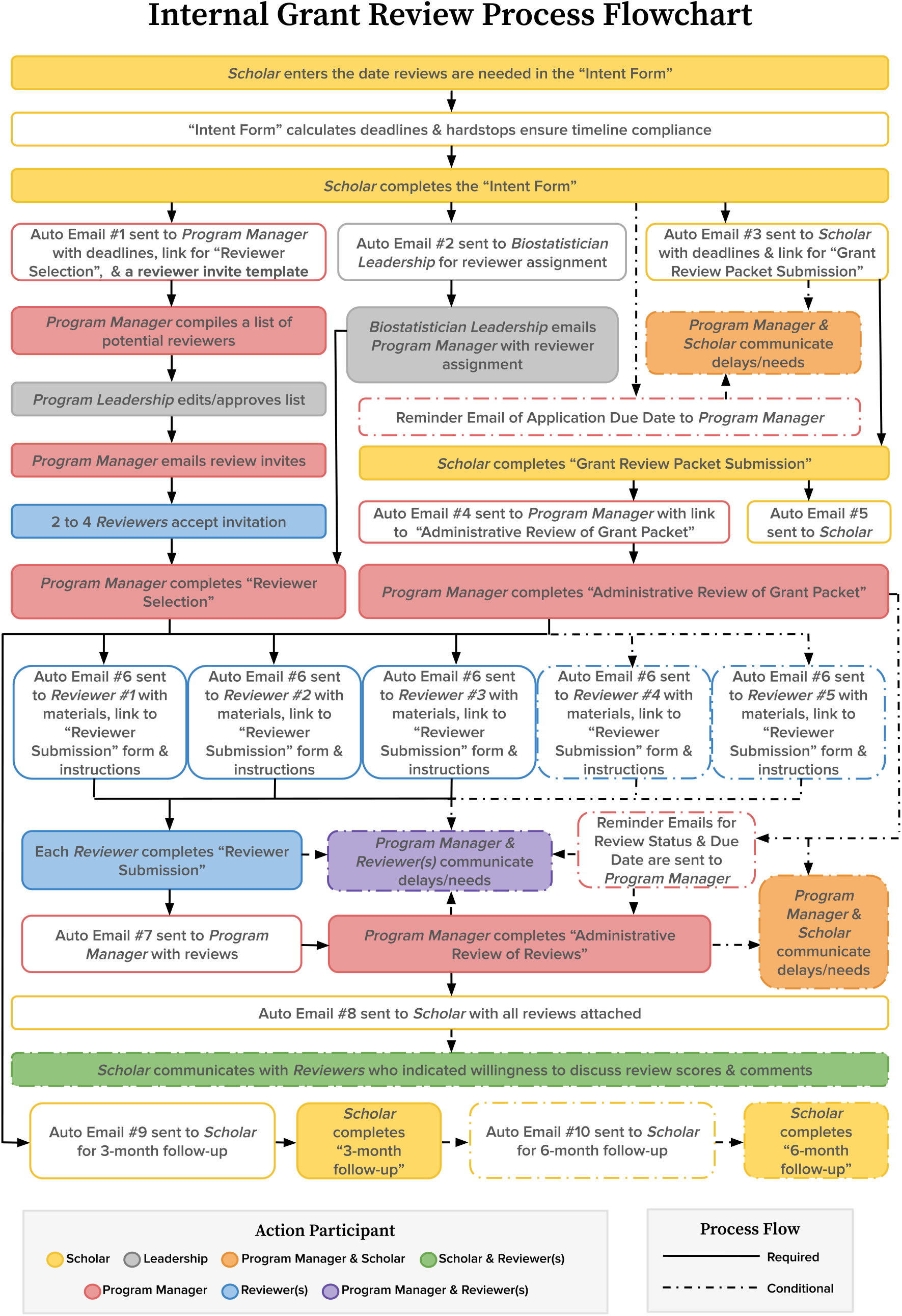
Process Flowchart for the Internal Grant Review Program.

### Evolution of Internal Grant Review Program

The IGR program was first piloted for three consecutive iterations: winter 2020, spring 2021, and summer 2021. Anecdotal positive feedback from Scholars encouraged the iTSP Leadership to continue to offer the resource; however, the management of the IGR program at the time required more manual effort from the iTSP Leadership than was sustainable. The program’s process required improvement to allow for multiple grant application submissions and year-round offering. REDCap was identified as a viable solution to this process problem, providing the ability to automate and track the IGR program’s workflow, collect and secure data, and generate regular reports for the iTSP Leadership (see Figure 1).

The evolution of the IGR program workflow has resulted in greater efficiency; data collection, progress tracking, communication, task reminders, and reporting are all enhanced by the use of automation in REDCap. Overall, the IGR program process time has been reduced from eight weeks to six weeks. Less labor effort is required from the iTSP Leadership members with our latest iteration, directly resulting in the year-round offering of this resource. Scholars benefit from the improvements to the program process with more time to prepare their application submissions for internal review without sacrificing the time in which they receive their reviews. Data collection is easier to obtain from all parties involved with the use of surveys, forms, and automated alerts and notifications. Follow-up data on application results is being routinely collected as well via scheduled automated survey invitation emails at three and six months after the Scholar has participated in IGR. Tracking and program evaluation are significantly simpler as well due to REDCap’s dashboard, reporting, and exporting features. Additionally, because REDCap is available at no cost to nonprofit organizations and offers project file-sharing capabilities, our IGR can be adapted for other programs and institutions. See Supplementary Materials for the project files.

### Program Evaluation

#### Program Participation and Grant Statistics

The IGR program is in its fourth year of being offered. Internal reviews have been conducted for 26 of the 39 Scholars eligible to participate. In total, 29 proposals have gone through our internal review program (14 K proposals and 15 R proposals) prior to NIH submission. Four of these proposals were submitted to our program for a second internal review, making the total number of applications internally reviewed 33. Out of the 29 proposals reviewed in this program, 28% are being actively pursued by Scholars with funding results pending. Excluding proposals still awaiting funding decisions, 43% of the proposals (9 out of 21) submitted to IGR ultimately secured funding, either from the NIH (8 proposals) or a foundation award (1 proposal). Excluding unfunded Scholars with proposals still pending, 75% of Scholars (15 out of 20) who have utilized IGR have achieved independent funding, either with the proposal submitted to IGR or a subsequent one.

#### Scholar Feedback

Scholars often report to the iTSP Leadership and their Scholar peers satisfaction with IGR. In October 2023 a REDCap feedback survey about the program was sent via email to all 20 Scholars who had participated in IGR at the time. We had a 60% response rate to the anonymous survey. Of the Scholars who responded to the survey, 92% stated that the IGR program was “helpful”, “useful”, and/or “valuable”. One Scholar reported “This is an amazing resource that everyone should use! The feedback I received was indispensable and significantly strengthened my grant application”. Others said “It was extremely helpful. I think it helped my grant get funded”, “It was accurate and valuable feedback. I intend to use this review process for future grants”, and “The comments by reviewers who were subject matter experts really pushed the application to a fundable place”. While nearly all of feedback was positive, one Scholar expressed disappointment and concerns about perceived appropriateness of a specific reviewer’s qualifications.

Since the initiation of IGR, recurring comments from several Scholars who had participated in IGR have been observed by the iTSP Leadership (see Table 1.). To evaluate if these comments were common among IGR participants, a section was included in the survey listing them as potential “lessons learned”. Scholars were asked to select all statements that apply to their experience with IGR. Almost all (92%) said they learned that more time was needed than they had thought to prepare the application and 33% said they wished they’d submitted a near final draft for review. One Scholar even added the comment “It felt much more like an actual review that would have been more appropriate for a finished product (which mine was not). If I went through it again I would submit something closer to a final draft”. Only one Scholar reported they learned they needed more time to incorporate their reviewer feedback.

**Table 1.**
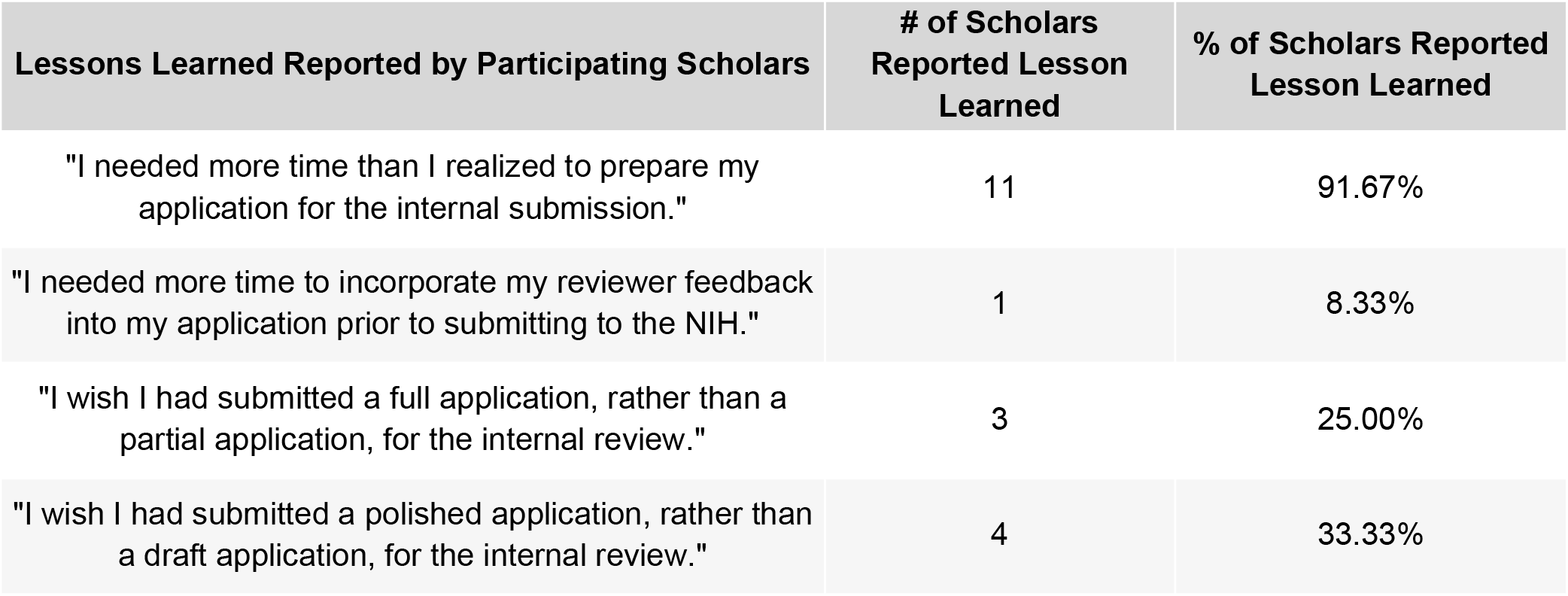
Results of “Lessons Learned” from Scholar participant feedback survey.

| Lessons Learned Reported by Participating Scholars | # of Scholars Reported Lesson Learned | % of Scholars Reported Lesson Learned |
| --- | --- | --- |
| "I needed more time than I realized to prepare my application for the internal submission." | 11 | 91.67% |
| "I needed more time to incorporate my reviewer feedback into my application prior to submitting to the NIH." | 1 | 8.33% |
| "I wish I had submitted a full application, rather than a partial application, for the internal review." | 3 | 25.00% |
| "I wish I had submitted a polished application, rather than a draft application, for the internal review." | 4 | 33.33% |

Net Promoter Scores (NPSs), a benchmark used to gauge customer satisfaction,^8,9^ were also collected in the Scholar feedback survey. In general, calculated NPSs above 0 are rated as “good”, above 20 are “favorable”, above 50 are “excellent”, and above 80 are “world-class”.^10^ Table 2. details our NPS question and individual scores. Based on the survey results, the NPS collected and calculated in October 2023 for the IGR program equaled 58, considered “excellent”, indicating that Scholars are overall satisfied with the program and are likely to recommend it to others.

**Table 2.** Net Promoter Score (NPS) results from Scholar participant feedback survey from October 2023.

| How likely are you to recommend using the Internal Grant Review to a Scholar peer?<br>(0=Not at all likely, 10=Extremely likely) |  |  |
| --- | --- | --- |
| Promoters | # of 10s | 6 |
|  | # of 9s | 2 |
| Passives | # of 8s | 3 |
|  | # of 7s | 0 |
| Detractors | # of 6s | 0 |
|  | # of 5s | 1 |
|  | # of 4s | 0 |
|  | # of 3s | 0 |
|  | # of 2s | 0 |
|  | # of 1s | 0 |
| Number of Responses |  | 12 |
| Calculated NPS Score |  | 58.33 |

#### Reviewer Statistics and Feedback

The IGR program has had 116 reviews total for the 33 applications submitted by Scholars. While reviewers are allowed to provide their critiques anonymously, more than half volunteer to meet with the Scholar afterwards, if the Scholar wishes, to provide additional context and support. In October 2023 we sent all reviewers who had participated in IGR a feedback survey and had a 36% response rate. Of the reviewers that provided anonymous feedback, nearly all (96%) reported they felt that they had scientific expertise aligned with the proposal they reviewed. Reviewers were asked to estimate the number of hours spent reviewing an application. The majority (42%) reported spending one to two hours on the application review. The next most reported amount of time spent was three to four hours (39%). Nearly all (96%) survey respondents indicated willingness to review again for future applications; most reporting willingness to review one per year. One reviewer stated “this is a very, very good program for aspiring scientists that are to be expected to get federal funding. You are helping by making the review process much like it would be for a NIH grant. Keep up the good work.”

Included in the reviewer feedback survey was a section soliciting advice from reviewers, of which 58% (15 out of 26) of respondents provided. There was a mixture of advice received, such as one that requested “more time to review” and another advised to make sure the reviewers “know the mechanism” for which they are reviewing. Two reviewers suggested for the final results of proposals to be reported to the reviewers. Three reviewers suggested that submitted proposals to the IGR program be “complete” and/or “fully developed” for a comprehensive evaluation.

## Discussion

The data for the Internal Grant Review program provides us with insight into how the program is being perceived and its impact on our Scholars. First, these results indicate that IGR is a resource that participating Scholars find to be of value and use. Second, participating in the IGR process can teach Scholars to better prepare their applications to maximize the benefit of a pre-review. Third, the limited data we have suggests that reviewers for the program are supportive and some prefer to see well-developed, polished proposals when reviewing.

Subjective feedback from Scholars (both anecdotal and solicited) has been consistently positive, indicating high satisfaction with IGR. Additionally, the NPS score for IGR provides quantifiable validation of this subjective feedback. The 43% success rate of proposals reviewed by IGR, higher than the reported 14% NIH success rate^2^, is another indicator of this resource’s usefulness. We recognize this is a relatively small sample size and further analysis is needed as more internal reviews are performed and funding results for Scholar’s proposals become available.

Feedback from both Scholars and reviewers suggests a need for purposeful messaging to Scholars to increase their awareness of allocating appropriate time for application preparation and optimizing the benefit of IGR by submitting a near complete application. In addition to updating the written IGR guidance with this information, the iTSP Leadership has started initiating early conversations with Scholars who are targeting NIH application cycles, ensuring they are aware of the “lessons learned” reported by past IGR participants. Additionally, to help Scholars with planning out their application preparation timeline, a tool for calculating due dates was created in Google Sheets for Scholars to use (see Figure 2.).

**Figure 2.**
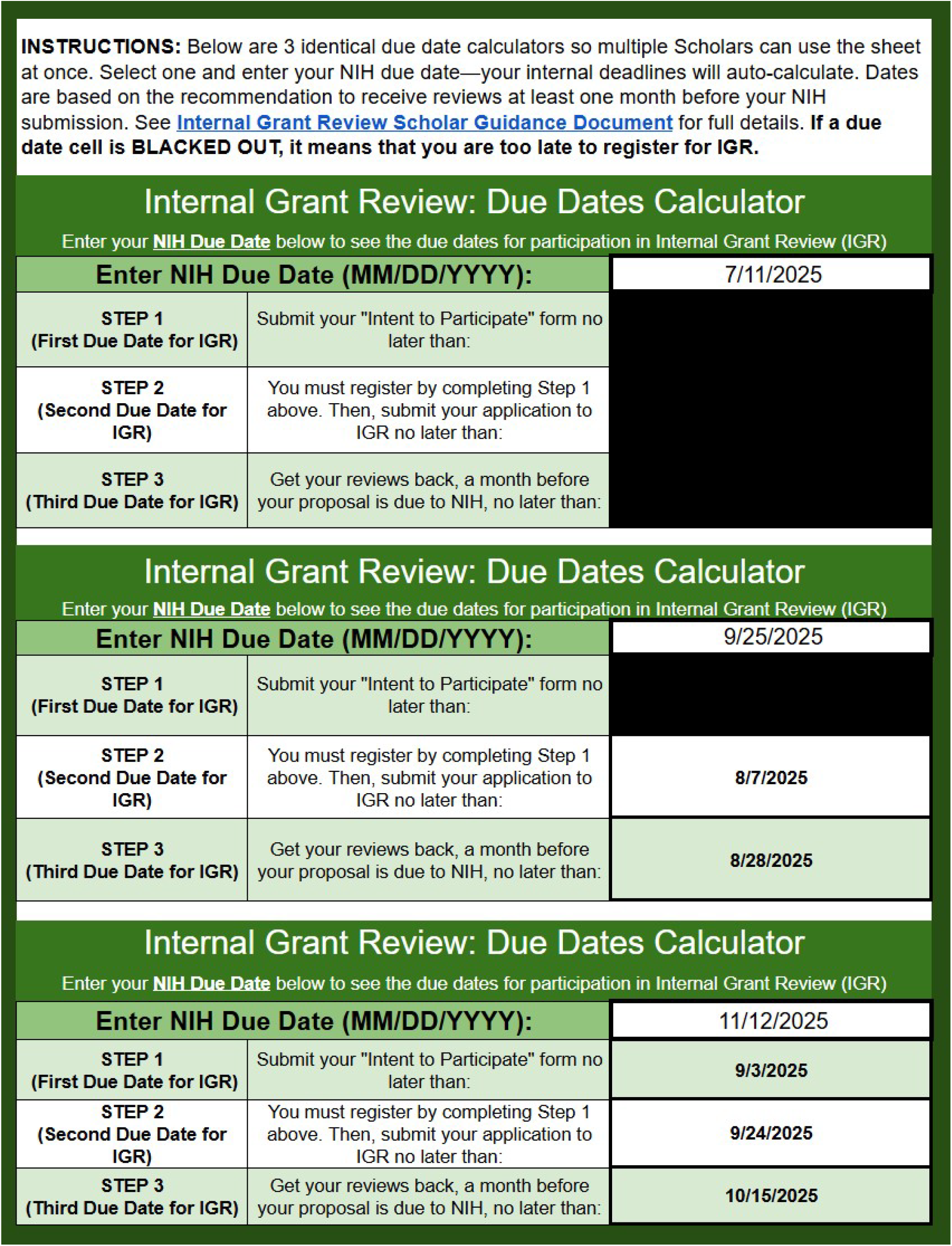
Date Calculator. created in Google Sheets for Scholars to utilize to determine IGR deadlines.

In response to the reviewer feedback we have already received, a new procedure will be implemented to express our appreciation through an annual email to all reviewers to include a report of the Scholars who utilized IGR and received funding. Some suggestions from Scholars and reviewers, such as providing more time for reviewing (reviewer) and a more rapid review of major proposal components (Scholar), are noted as potential areas of refinement, but will not be acted upon until further data warrants them a necessity. Future iterations of the IGR program will incorporate our feedback survey into the IGR process workflow so that any Scholar and reviewer who participates in IGR will be given the opportunity to provide anonymous feedback, including an NPS rating. In this feedback survey we will also include new data points to help us compare Scholar experiences with the real NIH review process to ours. These changes will enhance routine program evaluation that identifies IGR strengths and weaknesses.

## Conclusion

The Internal Grant Review program is a valuable offering for our early-career faculty appointed to the iTHRIV Scholars Program. As one Scholar stated in survey feedback, they “are lucky to have this resource available”. With iterations of the program, we’ve enhanced our offering by incorporating changes based on feedback and utilizing REDCap, making the program an efficient, sustainable, and constant resource. After several years of offering the program, it is evident that IGR has been successful in not only exposing the Scholars to an NIH-like review of their applications, but also providing the opportunity to go beyond their network to get expert guidance to strengthen their proposals prior to NIH submission. Our approach using the automated REDCap system may be useful to other programs offering grant review prior to grant submission.

## Supporting information

IGR REDCap Project Files

## Funding

The work in this publication was supported in part by the National Center for Advancing Translational Sciences of the National Institutes of Health under Award Numbers KL2TR003016 and UL1TR003015. The content is solely the responsibility of the authors and does not necessarily represent the official views of the National Institutes of Health.

## Conflicts of Interests

None declared.

## Notes

### Competing Interest Statement

The authors have declared no competing interest.

